# Antigenic escape accelerated by the presence of immunocompromised hosts

**DOI:** 10.1101/2022.06.13.495792

**Authors:** Ryuichi Kumata, Akira Sasaki

## Abstract

The repeated emergence of SARS-CoV-2 escape mutants from host immunity has obstructed the containment of the current pandemic and poses a serious threat to humanity. Prolonged infection in immunocompromised patients has received increasing attention as a driver of immune escape, and accumulating evidence suggests that viral genomic diversity and emergence of immune-escape mutants are promoted in immunocompromised patients. However, because immunocompromised patients comprise a small proportion of the host population, whether they have a significant impact on antigenic evolution at the population level is unknown. We used an evolutionary epidemiological model combining antigenic evolution and epidemiological dynamics in host populations with heterogeneity in immune competency to determine the impact of immunocompromised patients on the pathogen evolutionary dynamics of antigenic escape from host immunity. We derived analytical formulae of the speed of antigenic evolution in heterogeneous host populations and found that even a small number of immunocompromised hosts in the population significantly accelerates antigenic evolution. Our results demonstrate that immunocompromised hosts play a key role in viral adaptation at the population level and emphasize the importance of critical care and surveillance of immunocompromised hosts.

## Introduction

The repeated emergence of new variants hinders the control of epidemics (Gandon et al., 2016; Koelle et al., 2022; Tao et al., 2021). By acquiring mutations to escape existing immunity, pathogens can continue to thrive in host populations. The current pandemic of SARS-CoV-2 is due to the repeated emergence and spread of variants with undesirable phenotypes, including increased infectivity and high mortality, known as VOCs (Variants of Concerns) (Boehm et al., 2021; Tao et al., 2021; Thye et al., 2021). Each time an existing strain is replaced by a VOC, the number of cases and deaths increases leaving a significant impact on society in terms of both public health and the economy. Accumulating evidence indicates that each VOC evades host immunity developed by previous infections with pre-existing strains and vaccines (Garcia-Beltran et al., 2021; Liu et al., 2022) and that multiple mutations in the S-gene region targeted by host immunity contribute to immune escape (Harvey et al., 2021). Thus, the emergence of new variants is undoubtedly an important factor in determining the dynamics of infectious diseases.

Prolonged and persistent infection may facilitate the emergence of variants of SARS-CoV-2 (Dennehy et al., 2022; Thng et al., 2021). Because mutations occur as pathogens replicate in the host body, the emergence of a new variant depends on the within-host dynamics. Pathogens that successfully infect the host are usually quickly eliminated by the immune system of the host. However, in hosts with compromised immune function due to medical treatment or diseases such as AIDS, elimination of pathogens may be delayed, resulting in prolonged infection. Recently, long-term infection with SARS-CoV-2 in immunocompromised patients has been reported (Corey et al., 2021; Khatamzas et al., 2021; Leung et al., 2022; Martins-Chaves et al., 2020). The longer a pathogen stays in an infected person, the more it replicates, increasing the probability of mutation. Thus, persistent infection likely influences the emergence of new mutations. Indeed, in the case of SARS-CoV-2, genomic diversity has been reported to result from increased mutations within immunocompromised hosts (Weigang et al., 2021), suggesting that an immunocompromised host may increase the likelihood of mutation. Therefore, host heterogeneity in the time from infection to recovery may affect the accumulation of undesirable mutations.

Although evidence for the promotion of mutations in immunocompromised hosts is accumulating, the extent to which the presence of immunocompromised hosts affects antigenic escape on a population scale remains unclear. Moreover, immunocompromised patients are rare, and whether the small number of such patients has a significant impact on viral immune escape at the population level has yet to be elucidated. In this study, we theoretically examine the evolutionary dynamics of pathogen antigenic escape in a host population with heterogeneity in recovery rates from infection to determine how such heterogeneity affects epi-evolutionary dynamics at the population level.

Several theoretical studies have attempted to elucidate the complex evolutionary epidemiological dynamics in which pathogens continue to evade host immunity by repeatedly changing their antigenicity (Gog and Grenfell, 2002; Haraguchi and Sasaki, 1997; Marchi et al., 2021; Sasaki, 1994; Sasaki et al., 2022; Sasaki and Haraguchi, 2000). These studies have revealed the speed of antigenic escape (Haraguchi and Sasaki, 1997; Marchi et al., 2021; Sasaki, 1994) and predicted cross-immunity-driven discontinuous outbreaks in both time and antigenic space (Gog and Grenfell, 2002; Haraguchi and Sasaki, 1997; Sasaki et al., 2022). They also evaluated the joint evolution of pathogen virulence and antigenic escape (Sasaki et al., 2022). However, few studies have examined the epidemiological dynamics of antigenic escape when heterogeneity in host traits, such as recovery rates, is present. Smith and Ashby’s stochastic simulations suggest that the presence of hosts with low recovery rates increases the l Composition of immunocompromised and immune-competent hosts in the infected ikelihood of overcoming the valley of the fitness landscape, but the impact of heterogeneity in host recovery rates on antigenic escape rates and epidemics in more general situations remains unresolved (Smith and Ashby, 2022).

This study aims to examine the impact of host heterogeneity in the recovery rate on the population-level dynamics of antigenic escape of pathogens. We construct a simple mathematical model that accounts for host heterogeneity and investigate the effect of heterogeneity on the evolutionary-epidemiological dynamics of pathogen antigenicity. We then study how the rate of antigenic escape, i.e., the rate of the accumulation of antigen-changing mutations in a pathogen, is affected by host heterogeneity. We also examine how epidemic control measures affect the speed of antigenic escape and discuss implications for efficient control measures to slow the accumulation of immune-escape mutants.

## Model

We consider a host population made up of two classes of individuals differing in their recovery rates from an infectious disease (Fig. 1A). Specifically, if we denote the immunocompromised hosts as class 1 and the remaining individuals as class 0, the recovery rate of class 1 is significantly smaller than that of class 0 (or the mean infectious period is significantly longer in class 1 than in class 0). We consider the epidemiology and evolution of antigenic variants in such a heterogeneous host population under the assumption that antigenic phenotypes can be indexed in a one-dimensional space ().

**Figure 1.**
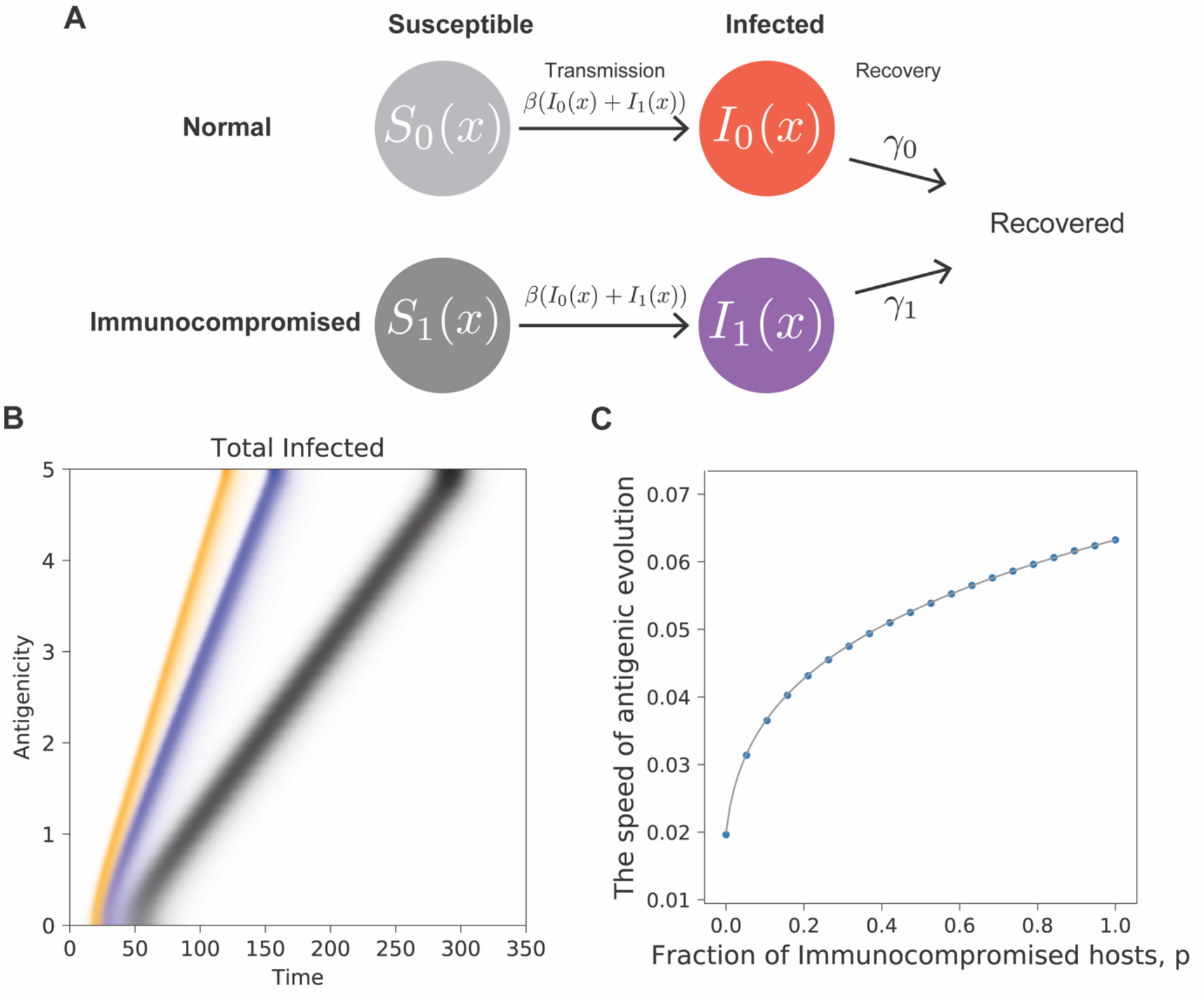
Antigenic drift in the population with heterogeneity in immune capacity. A. A graphical representation of the epidemiological model in the presence of immunocompromised hosts. B. The time changes the distribution of antigenicity obtained by numerical simulations of the model (1) with different proportions of immunocompromised hosts (shown by different colors). Antigenic drift in the population consisting only of hosts with normal immune capacity (*p* = 0) is shown in black, while that of populations where 10% and 30% of hosts are immunocompromised is shown in blue and orange, respectively. C. The speed of antigenic evolution as a function of the fraction *p* of immunocompromised hosts. The dots show the speeds obtained by numerical simulations of the model (1), and the solid curve is the analytical prediction calculated from equation (3). Parameters are *β* = 1.1, *γ*_0_ = 1, *γ*_1_ = 0.1, and *D* = 0.001.

We denote *S*_*i*_(*t, x*)as the proportion of class *i* hosts that are susceptible to the antigenic variant *x* of the pathogen at time *t. I*_*i*_(*t, x*)represents the proportion of class *i* hosts per unit area that are infected by the antigenic variant *x* at time *t*. The class *i* hosts that are susceptible to the antigenic variant *x* become infected with the force of infection *β*(*I*_0_(*t, x*)+*I*_1_(*t, x*))per unit time, where *β* is the transmission rate of the pathogen. We assume that the pathogen variants have identical traits except for antigenicity. An epidemic from an antigenic variant of a pathogen will end as the proportion of hosts susceptible to the variant is decreased. However, an immune-escaping variant can be generated through mutation before the focal epidemic is completely faded out to start another epidemic from the new variant. Random generation of antigenic mutants can be introduced by the diffusion term below, with diffusion constant *D* in the dynamics (Gog and Grenfell, 2002; Haraguchi and Sasaki, 1997; Sasaki, 1994; Sasaki et al., 2022). The infected hosts in class *i* then recover with the recovery rate *γ*_*i*_. Combining these epi-evolutionary processes, the proportions *S*_0_(*t, x*), *S*_1_(*t, x*), *I*_0_(*t, x*), and *I*_1_(*t, x*)change with time as

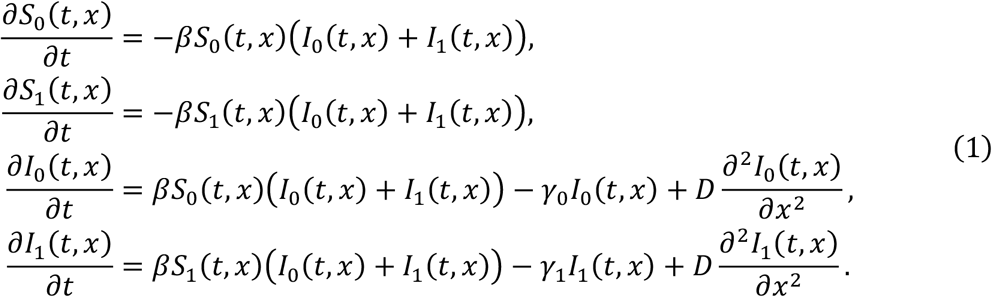

Though this is an SIR model (a compartment model for the changes in susceptible, infected, and recovered hosts densities), the proportions of the recovered or dead individuals are omitted as they do not affect the dynamics for *S*_*i*_ or *I*_*i*_. The diffusion coefficient *D* is calculated as the product of the mutation rate *μ* and the mean squared deviation of mutational change Δ*x* in antigenicity, *η*^2^ = *E*[(Δ*x*)^2^], using the equation *D* = *μη*^2^/2. The loss by recovery term in this model can be interpreted as any event responsible for the termination of infectiousness, including death caused by infection. In this paper, we assume that an immunocompromised host has an *m* times longer mean infectious period (time to recovery) than other hosts in the following equation:

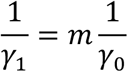

where *m* > 1. Another important parameter of the model is the fraction *p* of immunocompromised hosts in the population before the epidemic. Therefore, the initial proportion of hosts in each class at time *t* = 0 that are susceptible to a pathogen variant of antigenicity *x* is *S*_0_(0, *x*)= 1 − *p* and *S*_1_(0, *x*)= *p*, respectively. The present model (1) does not take into account the cross-immunity between different antigenic variants. The extended analysis of the model including cross-immunity is discussed in Supplementary Information.

Before showing the results of model (1), we briefly summarize how antigenic drift proceeds in a homogeneous host population where there is no host class 1 (*p* = 0) and all individuals have the same recovery rate *γ*_0_. The dynamics (1) are then reduced to:

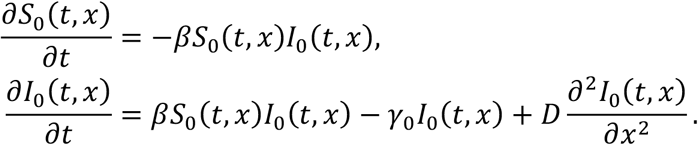

When a small amount of the pathogen variant with antigenicity *x* = 0 is introduced into the host population fully susceptible to the pathogen (*S*_0_(0, *x*)= 1 for all *x* and *I*_0_(0, *x*)= *ϵδ*(*x*), where *ϵ* is a small positive constant and *δ*(*x*)is the Dirac delta function), the pathogen persists by continuously evading host immunity as a traveling wave in antigenic space (Fisher, 1937; Gog and Grenfell, 2002; Haraguchi and Sasaki, 1997; Kolmogorov, 1937; Sasaki, 1994; Sasaki et al., 2022; Weinberger, 1982) with a constant wave speed:

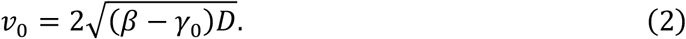

In other words, the speed of accumulation of antigenic mutants asymptotically approaches two times the geometric mean of the pathogen’s initial growth rate *β* − *γ*_0_ in a disease-free population and the diffusion constant *D* (half of the mutation variance). The total proportion of infected hosts maintained in the traveling wave ϕ = ∫ *I*_0_(*t, x*)*dx* approaches a constant determined from the following:

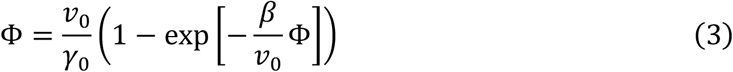

(Sasaki, 1994). Our aim in this paper is to extend these results to cases where the host population is heterogeneous to examine the effect on the speed of antigenic escape.

## Results

### Antigenic escape in a heterogeneous host population

The epi-evolutionary dynamics of antigenic escape of pathogens in a heterogeneous host population is studied using model (1). When a small number of hosts infected by a pathogen variant with a certain antigenicity (*x* = 0) is introduced into a disease-free host population, the primary outbreak occurs with the antigenic variant *x* = 0 providing that its basic reproduction number is greater than 1. Before the pathogen is cleared from the population from increased herd immunity, an antigenic mutant that can sufficiently escape host herd immunity raised by the primary outbreak could start the next outbreak. The sequence of such antigenic escapes by the pathogen and chase by host immunity could lead to the persistent epidemic of pathogens continuously changing their antigenicity (antigenic drift). The antigenic drift of pathogens can be modeled mathematically as coupled traveling waves of antigenic variants of the pathogen and antigen-specific immunity of a homogeneous host population (Gog and Grenfell, 2002; Haraguchi and Sasaki, 1997; Sasaki, 1994; Sasaki et al., 2022). Our model with host heterogeneity in immune competency showed stable coupled traveling waves of pathogen antigenic variants and specific host immunity in the antigenic space with a constant speed (Fig. 1B) as in the previous models (Gog and Grenfell, 2002; Haraguchi and Sasaki, 1997; Sasaki, 1994; Sasaki et al., 2022). However, we found that the wave speed, which corresponds to the speed of the accumulation of antigenic mutants, i.e., the pathogen’s speed of notorious evolutionary changes against host immunity and vaccination, sensitively depends on the host heterogeneity in immune competency (Fig. 1C), which is detailed in the rest of the paper.

If the width of the cross-immunity, which indicates the extent to which specific immunity raised at one antigenicity is effective against a similar antigenicity, is greater than the threshold, the coupled traveling waves of pathogen antigenic variants and specific host immunity become destabilized and exhibit a periodic sequence of outbreaks in time and antigenic space (Gog and Grenfell, 2002; Haraguchi and Sasaki, 1997; Sasaki et al., 2022) (Supplementary Fig. 1). Such large shifts in antigenicity are also observed in our model with host heterogeneity in immune capacity, and the periods in time and antigenic space between adjacent outbreaks depend on the heterogeneity (Supplementary Fig. 1). While such discontinuous antigenic escape is phenomenologically and practically important, the speed of the coupled traveling waves, which is our focus in this paper, is determined by the linearized system at the front end of the traveling wave and does not change by adding cross-immunity.

### Speed of antigenic escape and sensitivity to host heterogeneity

The linear analysis of the system (1) at the frontal end of the traveling wave of antigenic variants found in the Methods section reveals that the wave speed *v* in the heterogeneous host population consisting of the fraction 1 − *p* of hosts with a normal recovery rate *γ*_0_ and the fraction *p* of hosts with a lower recovery rate *γ*_1_ is given explicitly as:

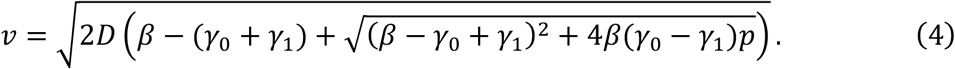

We assume that *β* > *γ*_0_ > *γ*_1_. This reduces to 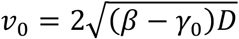, and to 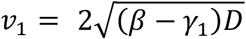, which match the expressions previously demonstrated to model the traveling waves in the homogeneous host population. The analytical formula (4) perfectly matches the wave speeds observed in the numerical simulation of (1) for varied *p* (Fig. 1C).

Equation (4) reveals several important results. First, the wave speed *v* of antigenic escape monotonically increases with the fraction *p* of immunocompromised hosts from *v* = *v*_0_ at *p* = 0 to *v* = *v*_1_ at *p* = 1. Second, the wave speed (4) is a saturating (or concave) function of *p*. Thus, the smaller the fraction of immunocompromised hosts, the larger its accelerating effect on the wave speed (Fig. 2). This has important implications for epidemic forecasting and its control, given the small proportion of immunocompromised hosts in most communities. Third, the closer the basic reproduction number *R*_0_ = *β*/*γ*_0_ of the pathogen in the population consisting only of hosts with normal immune capacity is to 1, the more sensitively the wave speed *v* increases with *p* (Fig. 2). This is most drastically shown with maximum sensitivity, (*dv*/*dp*)_*p*=0_, of the speed of antigenic escape to the proportion *p* of immunocompromised hosts at *p* = 0 (dashed lines in Fig. 2), calculated from (4):

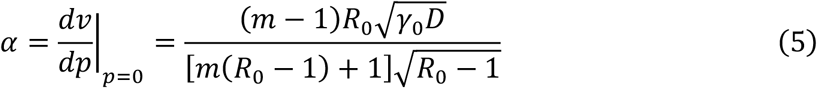

which diverges to infinity as the basic reproduction number *R*_0_ = *β*/*γ*_0_ approaches 1, where *m* = *γ*_0_/*γ*_1_ > 1 is the ratio of the infectious periods of immunocompromised hosts to that of hosts with normal immune competency. To summarize, the presence of immunocompromised hosts sensitively increases the speed of the accumulation of antigenic escape mutants, particularly when the proportion of immunocompromised hosts is small.

**Figure 2.**
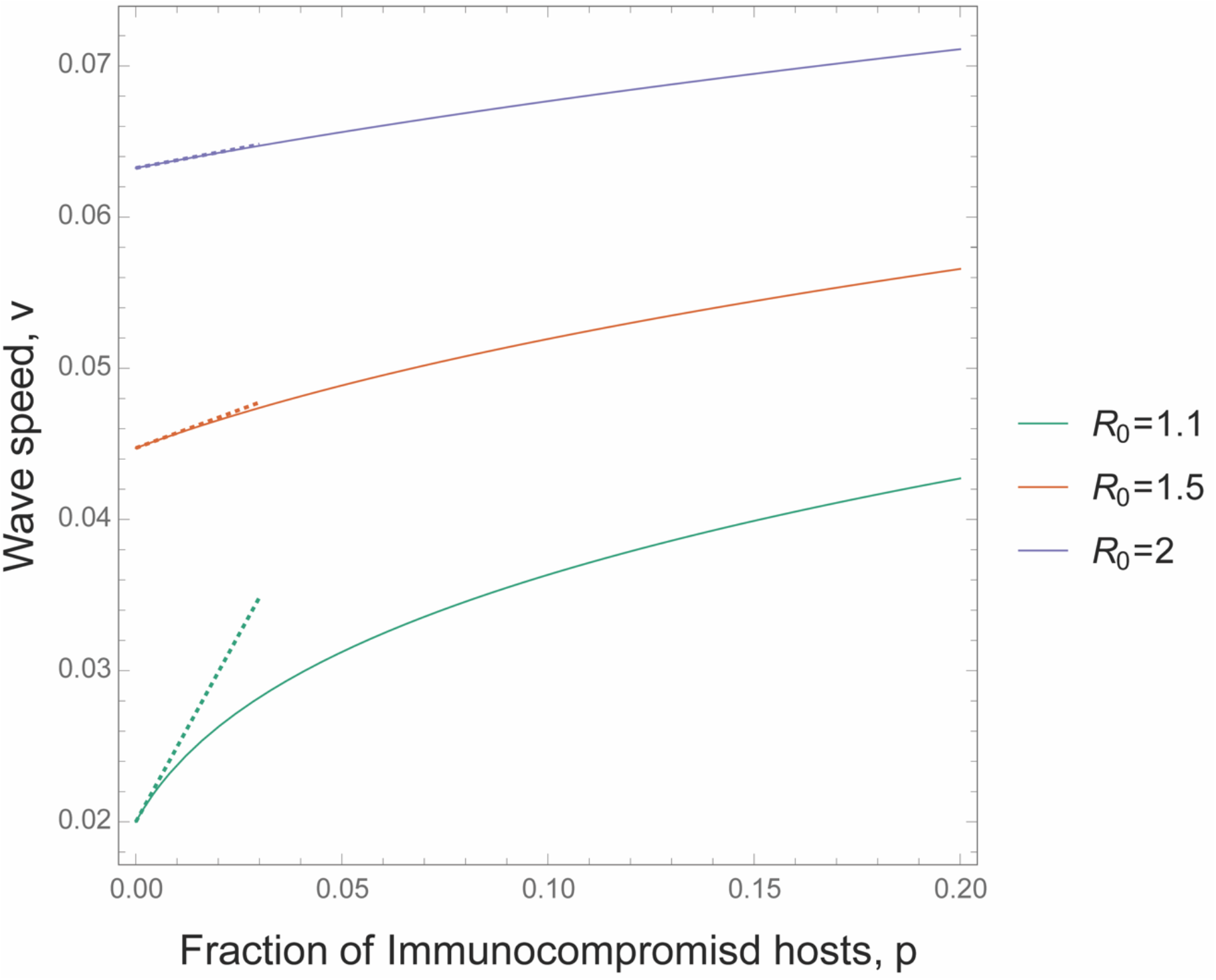
The sensitivity of wave speed *v* to the fraction of immunocompromised hosts. The solid curve is the wave speed calculated from (3). The dashed line is the slope of the curve at *p* = 0, which is *α* in (5). Different colors represent different *R*_0_ = *β*/*γ*_0_ values. The other parameters are *γ*_0_ = 1, *γ*_1_ = 0.1, and *D* = 0.001.

### Impact of immunocompromised hosts on disease dynamics

Each stable traveling wave of antigenic variants in host populations of heterogeneous immune competency (Fig. 1B) is decomposed into that in immune-competent hosts (blue) and immunocompromised hosts (red) (Fig. 3A). The asymptotic value of the total number of infected hosts at time *t*, 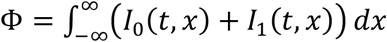, that quantifies the impact to the infectious disease caused by a continuously immune-evading pathogen, is determined from the following relationship, similar to (3) (see Methods for the deviation):

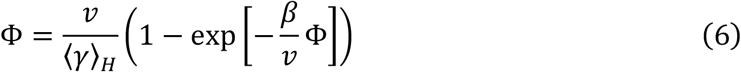

where *v* is the wave speed for a homogeneous host population defined in (4). Comparing this with (3) for a homogeneous host population, we see that the wave speed *v*_0_ in a homogeneous population is replaced by *v* in a heterogeneous population, and the recovery rate *γ*_0_ of immune-competent hosts is replaced by the harmonic mean of recovery rates, ⟨*γ*⟩_*H*_ = ((1 − *p*)/*γ*_0_ + *p*)/*γ*_1_)^−1^ in the heterogeneous host population (Fig. 3B). In addition, the ratio of infected immunocompromised hosts, 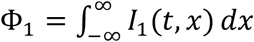 to that of immunocompetent hosts, 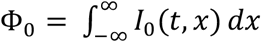, in a stationary traveling wave is shown as:

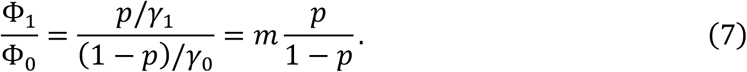

**Figure 3.**
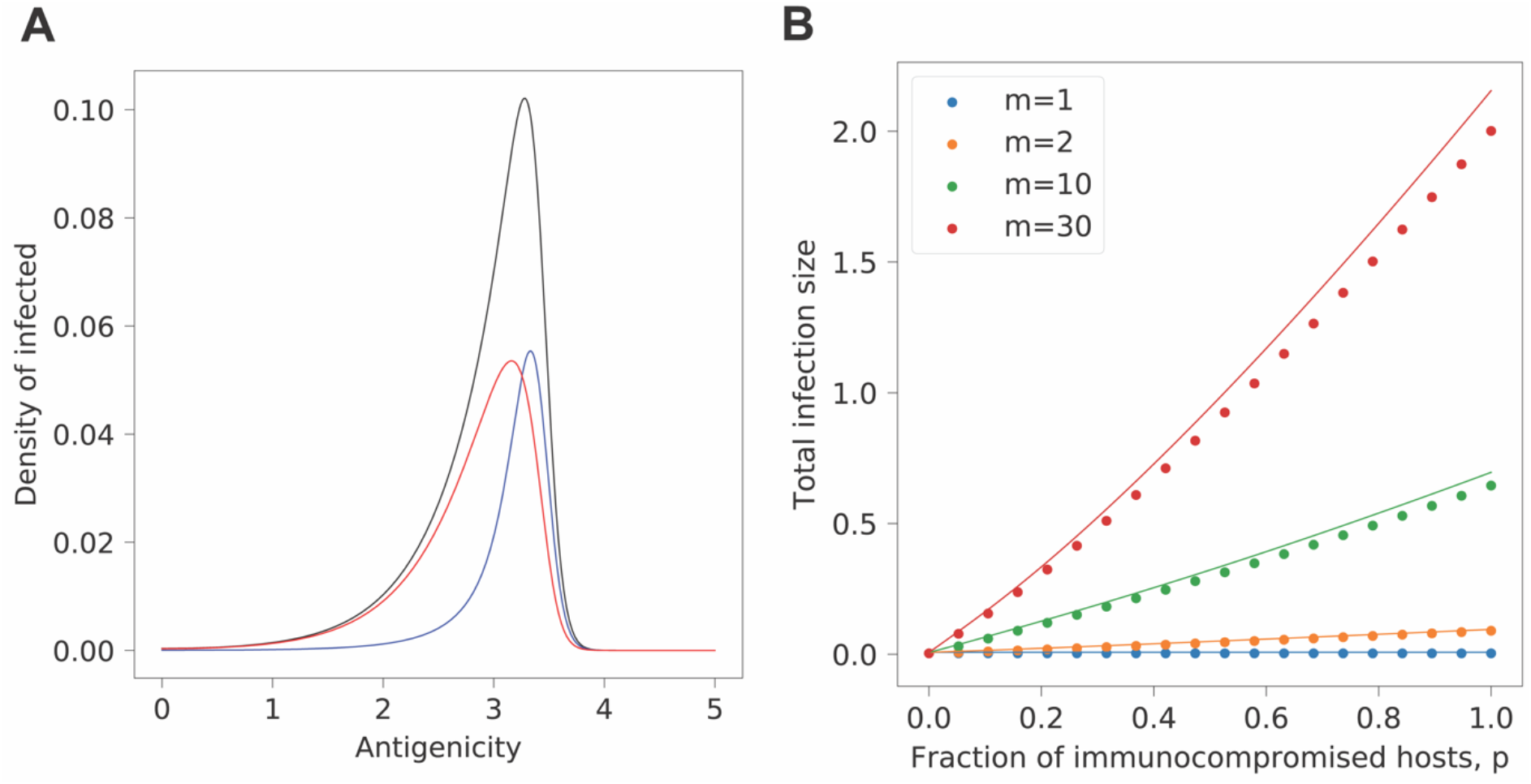
Composition of immunocompromised and immune-competent hosts in the infected population in a stable traveling wave in antigenic space. A. The traveling wave profiles of infected hosts over antigenic space in a stable traveling wave of antigenic escape. The black curve shows the total infected density, and the red and the blue curve represents the infected density of immunocompromised and immune-competent hosts, respectively. B. The total infection size Φ at a stable traveling wave is plotted against the fraction *p* of immunocompromised hosts. The dots are the results of numerical simulation of (1) and solid curves are the prediction from (6). Different colors are for different *m* = *γ*_0_/*γ*_1_ values. *m* = *γ*_0_/*γ*_1_ = 10 and *p* = 0.15 in A. The other parameters are *β* = 1.1, *γ*_0_ = 1, *D* = 0.001.

This implies that immunocompromised hosts are at *m* = *γ*_0_/*γ*_1_ times higher risk of infection than immune-competent hosts. Thus, not only do immunocompromised hosts disproportionately accelerate the speed of antigenic evolution of pathogens in the entire population, but they each bear a higher risk of infection than hosts with normal immune competency, suggesting the crucial importance of caring for immunocompromised hosts to protect the entire population and themselves.

### Control measures for suppressing antigenic escape

As an application of our model, we discuss intervention strategies to slow the evolution of antigenic escape. From a public health perspective, slowing the rate of antigenic change would be a great benefit. The slower rate of antigenic change increases the duration of time during which the existing vaccines are effective. Moreover, the longer time before immune escape increases the time available for the development of strain-nonspecific antiviral drugs. Thus, slowing the rate of antigenic change is vital for controlling persistent viral epidemics. As a strategy to slow antigenic escape, targeted interventions that concentrate countermeasures (e.g. vaccines, quarantine, etc.) on specific populations are considered (Fig. 4A). Here, intervention is assumed to reduce susceptibility. Let *σ*_*i*_ be the efficiency of the intervention on host class *i*, assuming that the susceptibility of the particular host population *i* is reduced to be 1 – *σ*_*i*_ times (*σ*_*i*_ ≤ 1) that before the intervention. The corresponding model for this case can be written as follows:

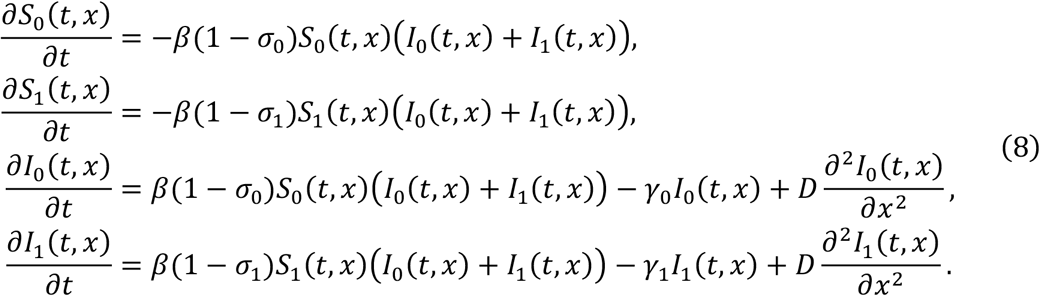

**Figure 4.**
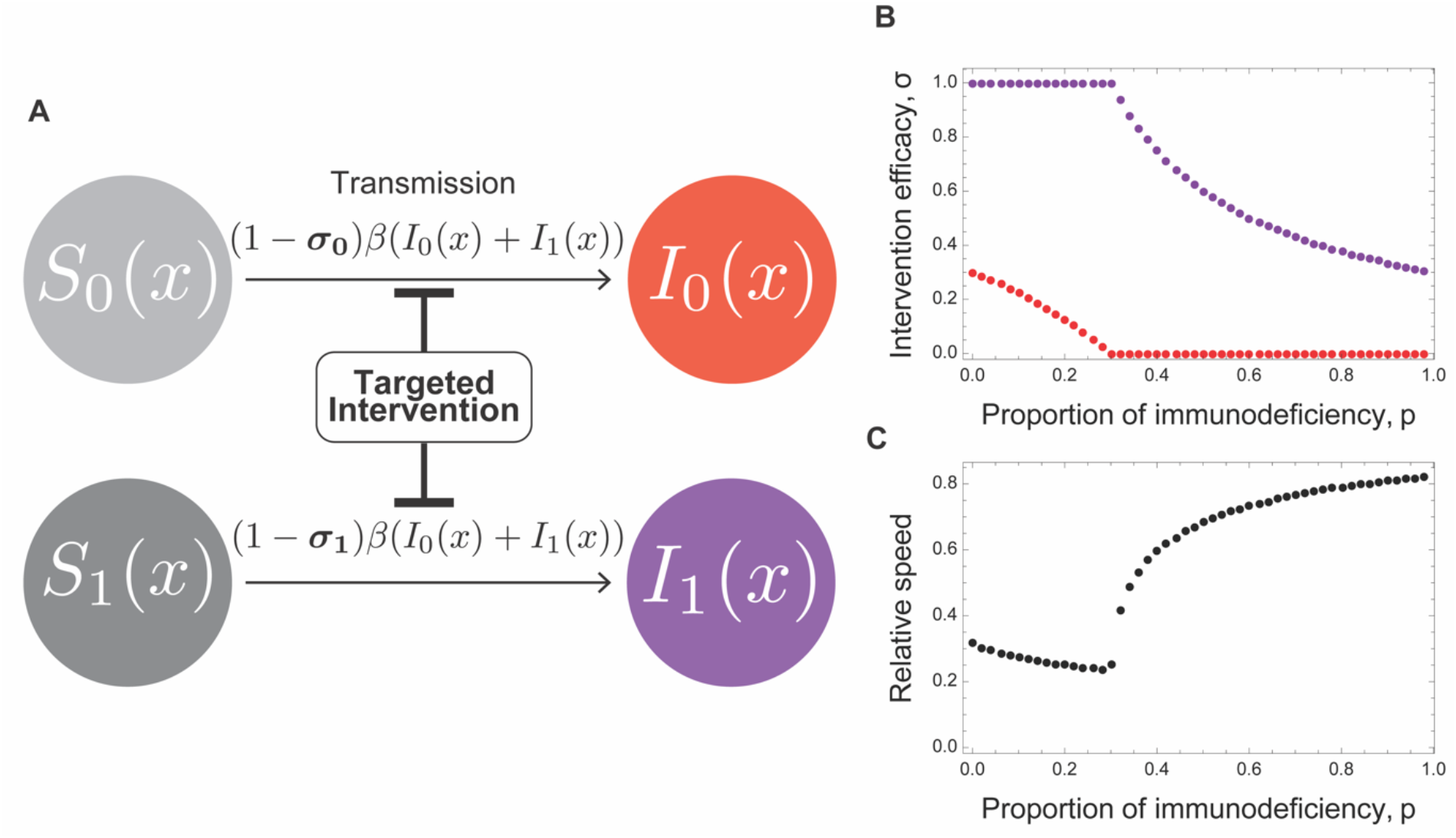
The optimal intervention scheme in the host population with heterogeneous immune competency. A. A graphical representation of the model of targeted intervention strategy. B. Under an economic cost-induced trade-off between *σ*_0_ and *σ*_1_, (1 − *p*)*σ*_0_ +*pσ*_1_ = *c*, the optimal intervention 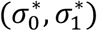 that minimizes the speed of antigenic escape evolution is shown for varied proportions of immunocompromised hosts. The purple and red dots show the optimal degrees, *σ*_0_ and *σ*_1_, of reducing susceptibilities of normal and immunocompromised hosts, respectively. C. The relative speed, *v*′/*v* under optimal intervention to that without intervention. The parameters are *β* = 1.5, *γ*_0_ = 1, *m* = 10, *D* = 0.001, and *c* = 0.3 (the economic budget for control is just enough to reduce the susceptibility of all individuals by 30%, or to perfectly reduce the susceptibility of 30% of the population, or a mixture of the efficiency and coverage).

This model contains heterogeneity in both recovery rate and susceptibility. We also calculate the rate of antigenic change in this model analytically as before (Methods), allowing us to examine how changes in susceptibility due to targeted intervention affect the rate of antigenic escape evolution.

A targeted intervention scenario is considered in which the intervention is applied to both immunocompromised and immune-competent hosts, but the degree of intervention varies between classes (Fig. 4B). The presence of immunocompromised hosts certainly accelerates the rate of antigenic escape evolution; however, because the proportion of immunocompromised hosts is usually smaller than that of immune-competent hosts, whether intervention should be focused on the small number of immunocompromised hosts or the large number of immune-competent hosts is unclear. We assume that the intervention has an economic cost *c*, which is given by the product of the degree of the suppression effect *σ*_*i*_ and the size of the target, 1 − *p* for *i* = 0 and *p* for *i* = 1:

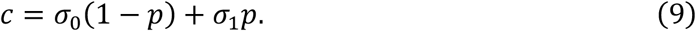

If we denote the speed of antigenic evolution under intervention by *v*′, the most efficient intervention for a fixed economical cost (9) is to minimize its ratio, *v*′/*v*, to the speed *v* before intervention. Thus, we calculated the optimal targeted intervention strategy, (*σ*_0_, *σ*_1_), that minimizes *v*′/*v* under the constraint (9) for varied proportions *p* of immunocompromised hosts (Fig. 4B). When *p* is less than the threshold *p*_*c*_ = *c* (the threshold is *p*_*c*_ = 0.3; Fig. 4B), the optimal intervention strategy 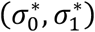 is to try to maximally reduce the susceptibility of immunocompromised hosts, while reducing the susceptibility of immune-competent hosts as far as the cost allows: 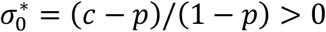 and 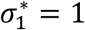 (Fig. 4B). In contrast, when *p* is greater than the threshold, the optimal intervention focuses only on reducing the susceptibility of immunocompromised hosts as far as the cost allows and completely ignore the intervention for immune-competent hosts: 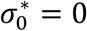 and 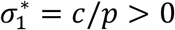 (Fig. 4B). These results show that the optimal intervention strategy prioritizes the intervention focusing on immunocompromised hosts, indicating that even if immunocompromised hosts exist only in small numbers, preferential intervention to them can efficiently reduce the rate of antigenic evolution. Interestingly, the relative wave speed *v*′/*v* under the optimal intervention control shows qualitatively different behaviors below and above *p* = *p*_*c*_ (Fig. 4C). Namely, if *p* is less than the threshold, the relative wave speed *v*′/*v* decreases with increasing *p*, indicating that the efficiency of the intervention increases with the number of immunocompromised hosts (Fig. 4C). In contrast, if *p* is above the threshold value, the speed of antigenic escape rapidly increases and approaches the value without intervention as *p* increases, indicating that the intervention rapidly loses its effectiveness as the number of immunocompromised hosts increases.

To be more practical, we need to carefully consider the more detailed relationship between targeted intervention measures, their effectiveness in reducing susceptibility, and their economic costs. However, this example illustrates the applicability of our model to design effective intervention policies to slow pathogenic immune escape.

## Discussion

The ongoing epidemic of SARS-CoV-2 has been accompanied by the emergence of several variants, and attention has focused on the evolutionary causes of such variants (Koelle et al., 2022; Tao et al., 2021). One mechanism that has been suggested is the presence of immunocompromised hosts in the population (Dennehy et al., 2022; Ghafari et al., 2022; Hill et al., 2022). In this study, we theoretically examine the impact of host heterogeneity in immune competency on the speed of antigenic escape of pathogens at the population level. Given that variation in immune competency is characterized by differences in recovery rates, we developed a model to represent the dynamics of antigenic drift in a population with heterogeneous recovery rates and derived the speed of the accumulation of antigenic escape mutants of a pathogen circulating in the population. We then examined the Composition of immunocompromised and immune effect of the presence of immunocompromised hosts on the accumulation of the antigenic escape mutants of the pathogen. The model reveals that the presence of immunocompromised hosts results in a faster accumulation of antigen-escaping mutations in the pathogen. In particular, we reveal that even if only a small proportion of immunocompromised hosts is present, the impact on the rate of antigenic evolution is significant. This emphasizes the importance of the presence of immunocompromised hosts not only on antigenic evolutionary dynamics at the individual level, but also at the population level. These results highlight the importance of careful surveillance and countermeasures for immunocompromised hosts (Li et al., 2022).

The presence of immunocompromised hosts not only sensitively accelerates the speed of antigenic escape of the pathogen, but also increases the magnitude of the pathogen prevalence in the entire population. Furthermore, immunocompromised hosts are to be found in the infected population at a much higher rate than in their original composition.

The strength of our study is that it provides an analytical and quantitative prediction of the extent to which the heterogeneity in host immune competency affects antigenic escape. To apply our model, data on the distribution of recovery rates in the host population is found to be particularly important, as we have revealed that the presence of hosts with longer infectious periods (lower recovery rates) could have a significant impact on the speed of antigenic evolution. Moreover, factors that may cause differences in recovery rates are not limited to immune competency. Other factors including age structure and vaccine dose condition affect host recovery rates. For example, older individuals recover more slowly than younger individuals (Voinsky et al., 2020), and unvaccinated individuals may experience slower recovery than vaccinated individuals (Chia et al., 2022; Singanayagam et al., 2022; Thompson et al., 2021). Thus, understanding the distribution of recovery rates shaped by these diverse factors is crucial.

We also applied our model to seek effective intervention strategies against pathogens capable of escaping host immunity. The targeted intervention to particular subpopulations is promising as any intervention including vaccination and behavioral regulation is accompanied by economic cost and overwhelming medical resources (Gandon and Lion, 2022; Gog et al., n.d.; Matrajt et al., 2021b, 2021a; Wang et al., 2021). Suppressing antigenic changes to slow the emergence of new immune-escaping variants is crucial because it allows time to develop vaccines against new variants and other countermeasures, such as antiviral drugs. In addition, slowing the rate of antigenic drift could lead to the containment of the epidemic with the existing vaccines. In considering effective measures, host heterogeneity in immune competency becomes again important. Our theory shows that effective measures in targeted interventions should focus on caring for hosts with weak immune competency (Fig. 4B). Because these hosts are at higher risk of infection than other hosts, a prioritized countermeasure protecting immunocompromised hosts has the advantage of protecting the entire population by slowing antigen escape. Because immunocompromised hosts are generally uncommon and specific interventions for such hosts are highly effective in slowing antigenic drift, priority measures targeted to them can be effective at relatively low economic and medical costs. While these results are based on several assumptions as to the effectiveness of targeted intervention and its cost, they indicate the potential to provide a basis for applying our model or an extended version of it to develop effective intervention strategies.

In a recent study, Smith and Ashby considered a similar model and investigated the effect of immunocompromised hosts on antigenic evolution (Smith and Ashby, 2022). Under the assumption that there is some correlation between pathogen infectivity and antigenicity, they numerically simulated their model and showed qualitative prediction that the presence of immunocompromised hosts allows pathogens to overcome an otherwise insurmountable fitness valley. Although our model is comparable to theirs except for their additional assumption on the correlation between antigenicity and pathogen fitness, our results give more general and analytical predictions on the rate of antigenic evolution.

Our theory can consider not only heterogeneity in recovery rates, but also more general heterogeneity, such as susceptibility and infectivity, and the rate of antigenic escape can be determined analytically (see Methods). Hosts have a variety of characteristics, including age, sex, and immune status. Heterogeneity in multiple host parameters, such as recovery rates and susceptibility, may be related to such host conditions. For example, heterogeneity in vaccination status creates heterogeneity in both susceptibility and recovery rates (Chia et al., 2022; Thompson et al., 2021). Recently, several studies have evaluated optimal vaccination policies to reduce the emergence of mutations that escape vaccines, taking into account host heterogeneity in the vaccination dose (Gandon and Lion, 2022; Gog and Grenfell, 2002; Saad-Roy et al., 2021). Our theory may also be useful in considering interventions to control vaccine escape mutants by considering vaccine-induced host heterogeneity. An interesting extension of our model for future research would be to determine how various intervention strategies affect the speed of antigen drift, while considering host heterogeneity.

Finally, we briefly situate this study within a theoretical context. In evolutionary ecology, much attention has been paid to the effects of heterogeneity on evolution in many contexts (Sparrow, 1999). This study theoretically addresses the speed of traveling wave solutions in heterogeneous populations. Although theoretical research on traveling waves is found in many evolutionary and ecological studies since Fisher’s work on the geographic spread of an organism (Fisher, 1937), the studies of traveling waves in heterogeneous populations remain scarce (but see Shigesada and Kawasaki, 1997). Thus, our theoretical approach holds promise for further development concerning directional evolution in heterogeneous populations.

## Methods

### Analytical calculation of wave speed in heterogeneous recovery host population

A traveling wave solution to the partial differential equations of our model (1) can be rewritten as a system of ordinary differential equations of *S*_*i*_(*z*)= *S*_*i*_(*t, x*)and *I*_*i*_(*z*)= *I*_*i*_(*t, x*) by introducing a moving coordinate with the same speed *v* as the traveling wave: *z* = *x* – *vt*:

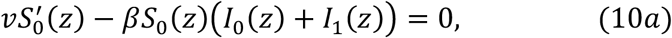

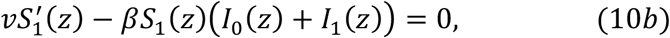

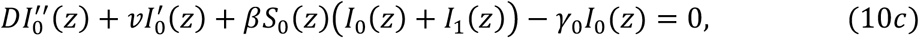

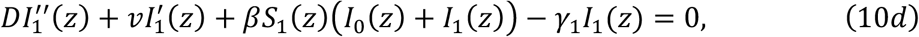

where ′ = *d*/*dz* and ′′ = *d*^2^/*dz*^2^ denotes the differentiations with respect to *z*. We linearize at the front end of the traveling wave of antigenic variants where all hosts are susceptible, *S*_0_(*z*)= 1 − *p* and *S*_1_(*z*)= *p*:

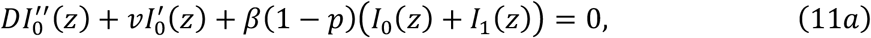

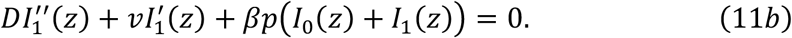

We then substitute exponential forms *I*_0_(*z*) = *ξ*_0_*e*^−*λz*^ and *I*_1_(*z*)= *ξ*_1_*e*^−*λz*^ where *λ* > 0 is the rate of exponential decay of traveling waves towards their frontal end and *ξ*_*i*_ > 0 is the positive constant representing the relative weights of immune-competent and immunocompromised hosts in the frontal end of the traveling wave. Substituting these into (11) yields:

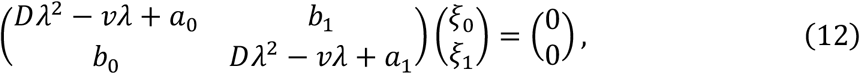

where *a*_*i*_ denotes the initial growth rate within class *i*, and *b*_*i*_ denotes the contribution of class *i* to the other class

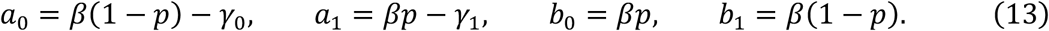

To obtain a nontrivial solution, (*ξ*_0_, *ξ*_1_)^T^ ≠ (0,0)^T^, to (12), the matrix in the left-hand side of (12) must be nonsingular, and hence, its determinant must vanish:

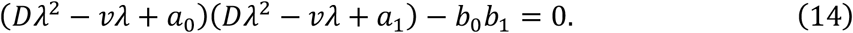

This is a quadratic equation for *𝒳* = *vλ* – *Dλ*^2^, *𝒳*^2^ − (*a*_0_ +*a*_1_)*𝒳* +*a*_0_*a*_1_ – *b*_0_*b*_1_ = 0, and is solved as:

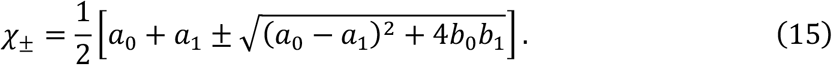

For the solution to be biologically feasible, both *ξ*_0_ and *ξ*_1_ must be positive, which means that only *𝜓*_*c*_ corresponds to a biologically feasible solution. Therefore, the relationship between the wave speed *v* and the exponential decay rate *λ* of the traveling wave towards the frontal end can be expressed as:

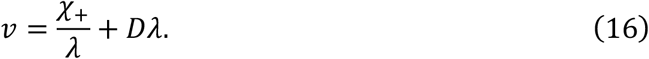

This is the dispersion relationship between the wave speed and the exponential decay rate of the traveling wave tail. In scalar equations, the traveling wave solution exists only for the wave speed *v* ≥ *v*_min_, where *v*_min_ is the minimum wave speed as a function of *λ*, and the stable wave solution for adequate initial distributions (e.g. the initial distribution is confined to the left of a certain position) has *v*_min_ as its speed. In a vector-valued system as in our model (which has a pair of distributions (*I*_0_(*z*), *I*_1_(*z*))), the same principle is mathematically proven under certain conditions (Fisher, 1937; Kolmogorov, 1937; Murray, 2002; Shigesada and Kawasaki, 1997); however, in some cases linearization of the system does not give the speed of the stable traveling wave (Hosono, 1998; Lewis et al., 2002). We have confirmed that the minimum wave speed in the linearized system (11) fits the speed observed in the numerical simulation of our model (1).

The wave speed *v*_min_ that minimizes (16) as a function of *λ* is:

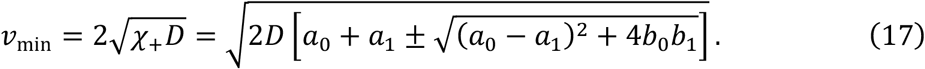

By substituting (13), we obtain analytical expression (4) in the main text:

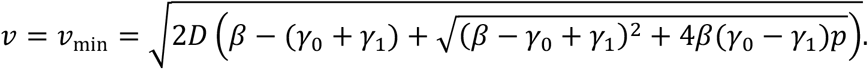

### Analytical calculation of total epidemic size and the ratio of infected immunocompromised hosts

In this section, we derive (6) and (7) in the main text. Integrating both sides of (10*a*) from *z* to ∞ give:

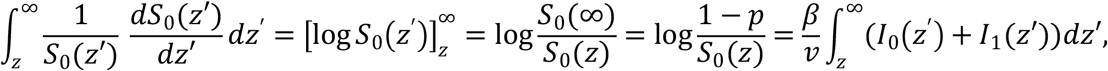

Where we used *S*_0_(∞)= 1 − *p*. This yields the following:

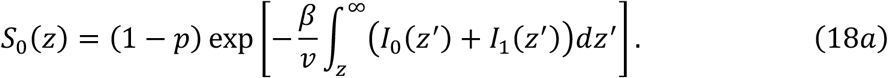

In the same vein, we obtain the following from (10b):

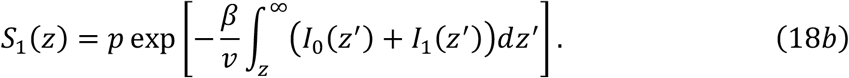

Substituting (18) into (10c) and (10d) to eliminate *S*_0_(*z*)and *S*_1_(*z*), and noting the following:

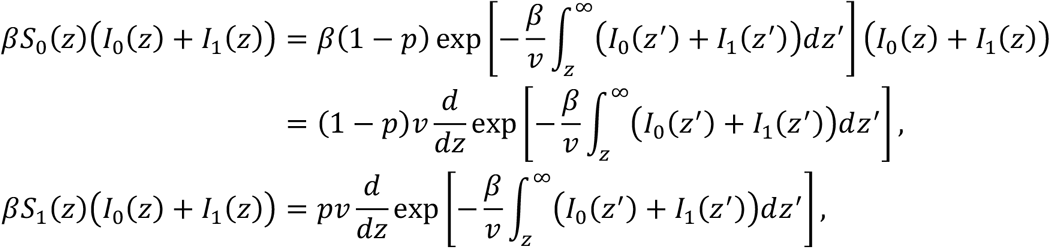

we obtain:

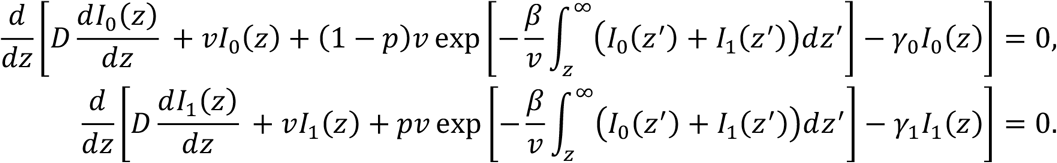

Integrating both sides from −∞ to ∞ and noting *I*_*i*_(±∞)= 0 and 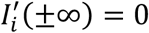, we then have:

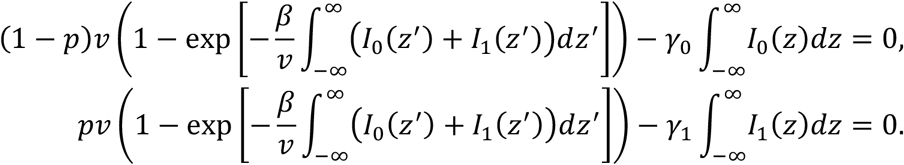

Thus, the following equations can be used for 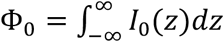 and 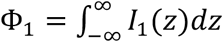:

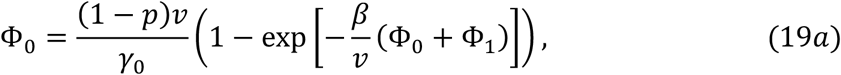

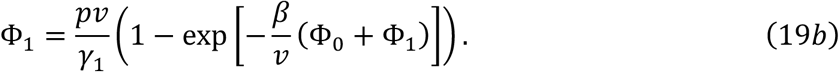

By taking the ratio of both sides of (19b) to those of (19a) we obtain (7) in the text:

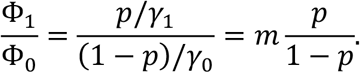

By adding (19a) and (19b), we obtain the equation (6) for the total epidemic size, ϕ = ϕ_0_ +ϕ_1_ in the main text:

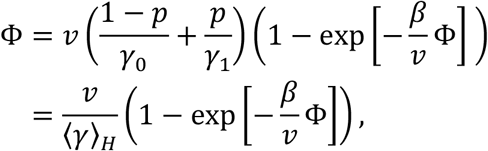

where ⟨*γ*⟩_*H*_= ((1 − *p*)/*γ*_0_+*p*/*γ*_1_)^−1^ represents the harmonic mean of recovery rates.

### Analytical calculation of wave speed in general heterogeneous host population

If heterogeneity is present in susceptibility and transmissibility as well as in recovery rates, model (1) is extended to the following:

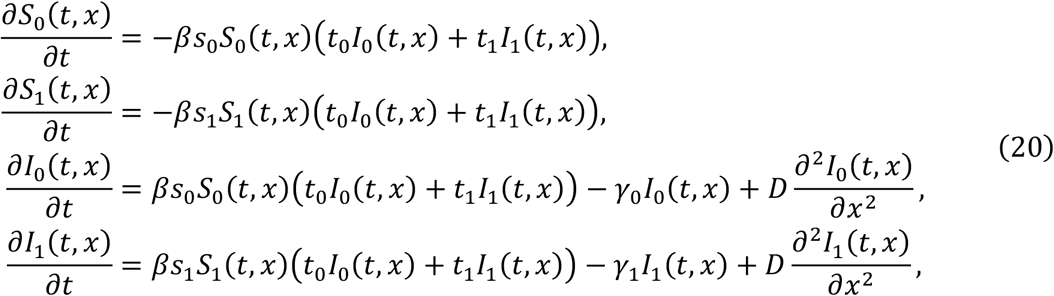

where *s*_*i*_, *t*_*i*_, and *γ*_*i*_ represent the susceptibility, transmissibility, and recovery rate of class *i* hosts, respectively.

We can obtain the analytical formulae in this model by using the above procedure. Briefly, linearizing this system at the frontal end of the wave, we get essentially the same equation as (12):

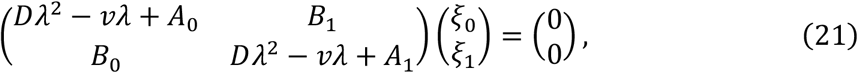

where *A*_0_ = *βs*_0_*t*_0_(1 − *p*)– *γ*_0_, *A*_1_ = *βs*_1_*t*_1_*p* – *γ*_1_, *B*_0_ = *βs*_1_*t*_0_*p*, and *B*_1_ = *βs*_0_*t*_1_(1 − *p*). We then obtain the formula for the wave speed:

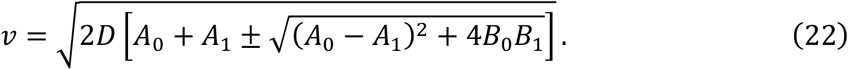

In the main text, we consider intervention strategies that target subpopulations in a specific manner. Intervention strategy (*σ*_0_, *σ*_1_)changes the susceptibility of class *i* hosts to 1 – *σ*_*i*_ times the susceptibility before intervention. In this situation, *A*_1_ = *β*(1 – *σ*_1_)(1 − *p*)– *γ*_1_, *A*_1_= *β*(1 – *σ*_1_)*p* – *γ*_1_, *B*_0_ = *β*(1 – *σ*_1_)*p, and B*_1_ = *β*(1 – *σ*_0_)(1 − *p*). Applying these to (22)gives the speed of antigenic evolution under the intervention strategies.

## Acknowledgements

We thank Hisashi Ohtsuki for helpful discussion.

## Competing interests

The authors declare no competing interests.

## Notes

### Competing Interest Statement

The authors have declared no competing interest.

