## Supplementalry Information for "Antigenic escape accelerated by the presence of immunocompromised hosts"

**Supplementary information for Antigenic escape accelerated by the presence of immunocompromised hosts**

**Cross immunity model**

We here extend our model (1) in the main body to include cross-immunity, as in Haraguchi and Sasaki (1997), Gok and Grenfell (20xx) and Sasaki et al. (2022). For simplicity, we assume the polarity immunity model introduced by Gok and Grenfell – a host infected by a pathogen of a particular antigenicity $y$ may acquire the perfect immunity against a different antigenicity $x$ $(\neq y)$ with probability $\sigma(x-y)$, but remain perfectly susceptible to antigenicity $x$ with probability $1-\sigma(x-y)$, where $\sigma(x-y)$ is a decreasing function of the distance $|x-y|$ between the antigenicities $x$ and $y$. With this assumption the dynamics with cross-immunity is

$$
\begin{aligned}
\frac{\partial S_0(t,x)}{\partial t} &= -\beta S_0(t,x)\int_{-\infty}^{\infty}\sigma(x-y)\big(I_0(t,y)+I_1(t,y)\big)dy\,, \\
\frac{\partial S_1(t,x)}{\partial t} &= -\beta S_1(t,x)\int_{-\infty}^{\infty}\sigma(x-y)\big(I_0(t,y)+I_1(t,y)\big)dy\,, \\
\frac{\partial I_0(t,x)}{\partial t} &= \beta S_0(t,x)I(t,x)-\gamma_0 I_0(t,x)+D\frac{\partial^2 I_0(t,x)}{\partial x^2}, \\
\frac{\partial I_1(t,x)}{\partial t} &= \beta S_1(t,x)I(t,x)-\gamma_1 I_1(t,x)+D\frac{\partial^2 I_1(t,x)}{\partial x^2},
\end{aligned}
\tag{$S1$}
$$

where $\sigma(d)=\exp(-d^2/2\omega^2)$ is assumed in the simulations below with the width of cross immunity $\omega$. In this model, the interaction between strains via cross-immunity results in the emergence of periodical outbreaks of the pathogen in both time and antigenicity space rather than continuous antigen drift (see Haraguchi and Sasaki 1997, Gok and Grenfell 20xx, Sasake et al. 2022). In Fig S1, these equations are numerically solved at the same parameter in Fig.1 in the main text except for the new parameter, the width of cross-immunity set to $\omega=1.2$. The linearized equation of $(S1)$ at the frontal end of the traveling wave is the same as that of the model in the main text, which determines the wave speed as is shown in Materials and Methods. Thus, the speed of antigenicity evolution in the cross-immunity model (S1) is the same as discussed in the main text.

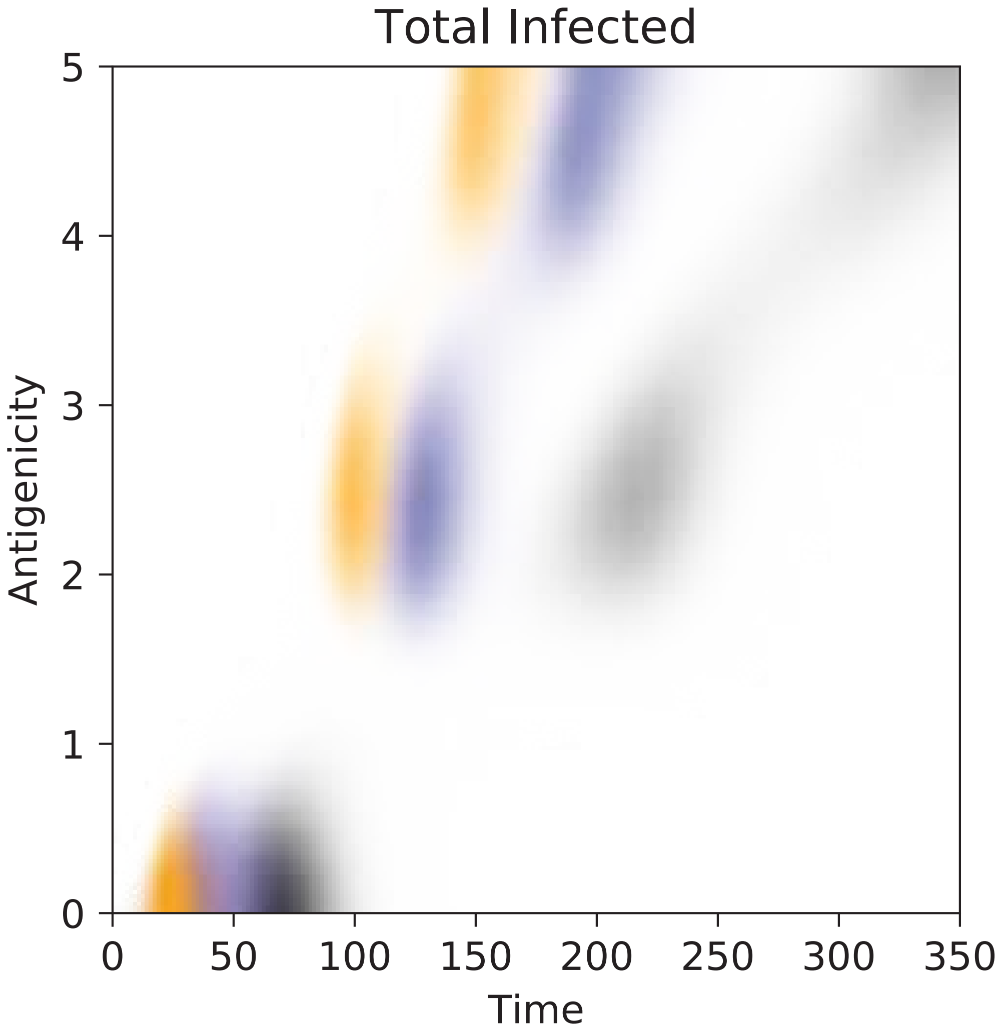


**Figure S1**. Antigenic shift in cross-immunity model. The time changes the distributions of pathogen antigenicity obtained by numerical simulations of the model (S1) with different proportions of immunocompromised hosts (shown in different colors). Cross-immunity kernels is $\sigma(x-y) = \exp(-(x-y)^2/\omega^2)$. The antigenic shift in the population consisting only of hosts with normal immune capacity ($p = 0$) is shown in black, while those in the population where 10% and 30% of hosts are immunocompromised are shown in blue, and orange, respectively. Parameter values are the same as in Fig.1 except for $\omega = 1.2$.
